# Comparative sensitivity of continuous cardiovascular markers for detecting cognitive workload, time-on-task fatigue, and acute stress

**DOI:** 10.64898/2026.09.02.749015

**Authors:** Pétur Kristófersson, Sigrún Þóra Sveinsdóttir, Sóley María Kristínardóttir, Stefnir Húni Kristjánsson, Kamilla Rún Jóhannsdóttir

## Abstract

Cardiovascular markers such as heart rate (HR), heart rate variability (HRV), and blood pressure (BP) are promising for low-intrusive monitoring of mental states relevant to safety-critical performance, yet evidence is mixed regarding which measures are most sensitive to cognitive workload, stress, and fatigue across different task types and time frames. Sixty adults (33 women; *M* age = 28.1 years, SD = 7.9) completed a 90-120 minute laboratory protocol including four cognitive workload tasks (Judgment of Line Angle and Position, Corsi Block-Tapping, Balloon Analogue Risk Task, Stroop), a modified Trier Social Stress Test (preparation, speech, arithmetic), and a fatigue condition operationalized as time-on-task during a prolonged Stroop (15 min) and Psychomotor Vigilance Task (10 min), each divided into three intervals. Continuous beat-to-beat HR, interbeat interval (IBI), and systolic/diastolic BP were recorded using the CareTaker device; HRV indices (RMSSD, high-frequency power) were derived from IBI using Kubios. Difference scores (task minus baseline) were analyzed using linear mixed models (random intercepts for participants; age and gender covariates), with Tukey-corrected pairwise comparisons to quantify sensitivity across conditions. HR and IBI consistently differentiated multiple workload and stress phases and distinguished Stroop-based fatigue from PVT intervals. HRV-RMSSD showed selective sensitivity, differentiating specific cognitive tasks (notably Stroop relative to other workload tasks), but only demonstrated a trend toward detecting fatigue-related differences at later time intervals. High-frequency HRV showed task-dependent effects only within the cognitive workload condition. Continuous BP did not reliably differentiate tasks, showing at most marginal (uncorrected) effects during TSST phase comparisons. Cardiovascular reactivity patterns depended on task type and temporal dynamics. HR and IBI provided the most robust sensitivity across workload, stress, and fatigue within a prolonged, multi-task protocol, while HRV indices contributed complementary but more task-specific information. The results support multi-metric monitoring approaches for real-time mental state transitions in high-risk operational settings.

## Introduction

In high-stakes occupations such as law enforcement, aviation, air traffic control, and emergency medicine, individuals are frequently exposed to significant stress, cognitive workload, and fatigue, increasing the risk of human error. Minimizing such errors is critical for ensuring safety and optimizing performance [1–5]. Prior research suggests that monitoring cardiovascular activity, including heart rate (HR), heart rate variability (HRV), and blood pressure (BP), may offer a cost-effective, low-intrusive tool for monitoring mental states, thus preventing situations where human error on task becomes almost inevitable [6–11]. Cardiovascular markers reflect immediate physiological responses [12] to cognitive and emotional challenges, making them suitable for detecting shifts in mental states due to increased task demands. However, despite their potential, inconsistencies remain regarding their sensitivity and reliability. While BP, HR, and HRV measures effectively differentiate extreme mental states (e.g., high vs. low workload), their ability to detect more subtle variations in stress, fatigue, or workload over time varies across studies [1,9,13–16]. These discrepancies may stem from common experimental limitations, such as inadequate task duration when assessing fatigue, a mental state that develops over extended time periods, or an over-reliance on single-task paradigms with varying difficulty levels [1,10,14,17–20], which fail to account for variability in cardiovascular responses across different task types or sustained performance demands. Additionally, a lack of standardization in task length and type further complicates cross-study comparisons, reducing the generalizability of findings across occupational settings [21,22]. To address these gaps, more research is needed to compare the sensitivity of multiple cardiovascular measures across diverse tasks, cognitive demands, and time frames. The present study aims to fill that gap by evaluating HR, HRV, and BP across varying mental states, including cognitive workload, acute stress, and time-on-task fatigue, using a complex and prolonged task setup. As such, this study seeks to identify distinct cardiovascular signatures associated with different cognitive challenges and determine which variables most effectively differentiate between mental state transitions. These findings will enhance the precision of non-invasive monitoring tools designed to improve safety and performance in high-risk professions.

### Prior work

#### Detecting cognitive workload

Both HR and HRV are widely regarded as useful indicators of cognitive workload. Numerous studies have shown workload-related increases in HR [8,23–33] and decreases in HRV [24,34–37] when comparing workload to non-workload or high workload to low workload conditions. A key challenge, however, lies in determining whether cardiovascular metrics can reliably track subtle, real-time variations in workload. Some studies suggest that HR does not consistently differentiate between multiple workload levels [3,38], whereas others demonstrate sensitivity to even small changes [14,28]. Recent reviews conclude that HR is among the more sensitive measures for capturing gradual shifts in workload, and meta-analytic evidence strongly supports the construct validity of both HR and HRV for workload assessment [21]. Still, their sensitivity may depend on task characteristics and duration.

Workload tasks relying heavily on working memory, such as mental arithmetic, appear to elicit stronger cardiovascular responses than those drawing on long-term memory or procedural knowledge [21]. HR may therefore be particularly responsive to tasks requiring sustained attention or immediate recall, making it well-suited to dynamic environments. Consistent with this, Raduntz et al. [39], looking at three-minute means taken early and late in a task that lasted for more than 20 minutes, found that HR is more sensitive to immediate workload changes, whereas HRV, particularly frequency-domain measures, reflects cumulative workload effects. Parent et al. [14] similarly found that HRV (RMSSD) distinguished between three levels of a modified N-back task, a paradigm in which workload accumulates over time.

Although HRV has been shown to distinguish between high workload and baseline conditions, its ability to capture smaller increments is less consistent [3,8,28]. Mehler et al. [27] reported that while HRV indices reliably detected difficult tasks, only about half differentiated moderate workload from baseline. In a simulated flight study with high overall demands, HR consistently tracked gradual increases in workload, whereas HRV metrics (high-and mid-frequency) were less reliable [40]. This suggests that HR may be more sensitive under high workload, while HRV may perform better in low-to-moderate ranges, though further testing is required. Similar findings were reported by Li et al. [41], who found that HR, but not HRV (HF/LF, SDNN, RMSSD), tracked load effects in multitasking flight tasks. Hidalgo-Muñoz et al. [8] also observed that HR, but not HRV, responded to changes in workload under high task demands. In contrast, studies using the interbeat interval (IBI), the fundamental beat-to-beat time series from which HRV indices, including RMSSD and HF, are calculated, have found the metric (which is an inversion of HR) to be sensitive to workload changes. For example, IBI successfully differentiated three levels of workload in Finnish Air Force pilots performing complex simulated flights [10]. In a study by Tjolleng et al. [18], IBI was more sensitive than HRV-RMSSD, being able to distinguish between three levels of task difficulty where participants performed on the N-back task while driving in a simulator. HRV-RMSSD could only differentiate between two difficulty conditions. Other HRV measurements were altered by increased cognitive workload but were insignificant. Wei et al. [42] showed reductions in RR interval variability with increasing task difficulty.

Blood pressure (BP) has also been linked to workload. Although Ayres et al. [21] could not conclude on BP, prior studies reported increases in systolic blood pressure (SBP) and diastolic blood pressure (DBP) with rising task demands [31,43,44]. For example, Veltman and Gaillard [45] found that SBP differentiated task levels in simulated flights, and DBP also reflected workload changes [38].

In sum, HR, HRV, and potentially BP provide sensitive indicators of cognitive workload, though their utility depends on task characteristics and workload intensity. HR shows consistent sensitivity across a broad range of demands, particularly in tasks requiring rapid recall or sustained attention, making it promising for real-time monitoring in dynamic environments [23,27]. HRV appears more effective in low-to-moderate workload ranges and may be better suited for detecting cumulative workload changes [39]. Studies also indicate that IBI may be sensitive for detecting workload changes. The temporal scale of the conditions being compared should be considered. When evaluating the sensitivity of cardiovascular variables for detecting momentary changes in mental states, it has been argued that shorter segments (30-120 sec) are better suited for capturing the actual impact a workload change may have on the cardiovascular system, at least for HR, BP and IBI [40,46]. At the shortest end, 30-40-second workload segments have been used for discriminative comparison [14,40]. More commonly, though, segments of two to three minutes are used in studies attempting to distinguish between increasing cognitive workload using cardiovascular variables [27,39,47]. Few studies compare workload periods of four or five minutes [3,8]. When it comes to HRV, recording periods of approximately five minutes are generally recommended for short-term HRV analysis [48]. However, other studies have suggested that even shorter recording periods may be sufficient for some HRV metrics [49].

#### Detecting stress

Cardiovascular measures such as HR, BP, and HRV have long been used to detect stress across various contexts [9,15,50–56]. Their sensitivity, however, appears to vary with the type and intensity of stressor as well as the timing of measurement [57].

In a study by Sequeira et al. [15], participants were exposed to three stressors of differing robustness (Trier Social Stress Task [TSST], karaoke singing, and unsolvable anagrams), with cardiovascular measures taken at baseline, before, after, and during recovery during five-minute segments. Both SBP and DBP distinguished baseline from stress reactivity during the TSST but not during the weaker stressors. In contrast, HR increased across all conditions, indicating greater sensitivity to less robust stress.

Hjortskov et al. [9] found that BP rose across introductory, stress, and control conditions during a computer-based task with psychological stressors. Even in the control condition, after stressors were removed, BP remained elevated relative to baseline, suggesting a slower return to resting levels. In contrast, HRV frequency metrics (HF, LF/HF ratio) distinguished stress from control conditions and detected recovery more effectively. These results suggest that BP is well-suited for capturing stress onset but less reliable for detecting rapid recovery, whereas HRV can track both onset and offset more dynamically. Pehlivanoğlu et al. [54] also reported that BP differentiated Stroop test difficulty but showed delayed recovery, reinforcing the idea that BP may lack sensitivity for immediate stress offset.

Overall, BP appears to respond reliably to robust stressors but less so to mild stress and may lag in detecting recovery [9,15]. HR, on the other hand, is effective for both mild and strong stressors, including unsolvable anagrams, Stroop tasks, IQ testing, and complex military simulations [15,20,55,56,58]. There is a growing consensus that HR and HRV outperform BP in detecting rapid, dynamic changes in stress onset and offset.

HRV has long been emphasized in stress research [51] because indices such as RMSSD indirectly reflect sympathetic nervous system activation through parasympathetic withdrawal [59,60]. This makes HRV an established marker of increased stress responses. Some researchers argue that HRV is preferable to HR because it directly reflects autonomic nervous system (ANS) activity [60]. However, findings from cognitive workload studies suggest that HR may be more robust in detecting subtle changes, while HRV can sometimes fail to capture small fluctuations in mental state changes [14,27]. In a study by Rajcani et al. [61], both HR and HRV (SDNN, RMSSD, and HF) showed a significant main effect of phases for a version of the TSST. In general, HRV decreased, and HR increased with stress (public speaking). No information was provided about follow-up comparisons of individual phases. In another study by Pulopulos et al. [62], HRV, measured as RMSSD, distinguished between baseline and a 15-minute anticipation period and then between the anticipation and a five-minute speech (modified TSST). Castaldo et al. [63] found that when using a three-minute mean, RMSSD did not decrease during a stress phase compared to a rest phase.

Recent advances in statistical modeling and machine learning have significantly improved HRV-based stress detection. Some studies combining multiple HRV variables report classification accuracies between 80–96% [64,65]. For example, Pourmohammadi and Maleki [66] achieved over 80% accuracy in distinguishing both three (no, moderate, high) and four (no, low, moderate, high) stress levels using HRV-derived features. These findings suggest that HRV, when supported by advanced analytic methods, may overcome some of its earlier limitations.

Stress detection tasks vary widely, from computer-based experiments [9] and social evaluation tasks such as singing or the TSST [15] to driving [16], combat simulations [55,56], and cognitive tasks like N-back, Stroop, and math problem solving [20,54,64,66,67]. Across these paradigms, both HR and HRV have consistently demonstrated sensitivity to stress of varying intensities. BP, in contrast, seems better suited to robust stressors and less reliable for detecting rapid or subtle stress.

In sum, HR, HRV, and BP all contribute valuable information for stress assessment, but their utility differs by stressor type and temporal dynamics. HR is broadly sensitive and easily measured in real-world settings, HRV is highly informative for autonomic responses and increasingly powerful when combined with modeling approaches, and BP remains most effective for robust stress but is limited in detecting quick recovery. Together, these measures provide complementary insights, with HR and HRV emerging as especially promising for real-time and dynamic stress monitoring. Studies quite commonly use three to five minute stress and no stress segments for comparison [57,63]. Some studies have used shorter segments such as 60 seconds [66].

#### Detecting fatigue

Cardiovascular measures show promise to detect fatigue, but findings have been somewhat mixed between studies. The focus here is primarily on task-induced fatigue [13]. Wright et al. [68] found that both SBP and DBP were more sensitive than HR in detecting fatigue. These results partly align with those of a study by Meng et al. [69], which found significant increases in SBP and DBP from pre-to post-fatigue measurements, but not HR. However, the results regarding HR in both these studies should be viewed with caution, since the literature generally shows that HR variables such as mean HR and HRV are generally sensitive indicators of fatigue [13,17,70]. Although Lu et al. [13], in their systematic review did not find a solid link between HRV and fatigue.

The results on HR and fatigue are generally mixed, with some studies showing that increased HR indicates fatigue [70,71], while others associate a decrease in HR with fatigue [13,17,72]. This could be due to the type of tasks performed, as some studies showing increased HR in association with fatigue are studies that use physical activity, such as combat simulation [70] and bicep curls mixed with physical work [71]. In contrast, the studies showing a decrease in HR with fatigue use tests such as the Gatekeeper task, a prolonged bimodal 2-back task [17]. A systematic review of 30 studies examining fatigue detection during simulated or real-world driving found that HR decreased across various fatigue-inducing tasks [13].

As with HR, a decrease or an increase in HRV during fatigue depends on the task type [13]. Prolonged tasks (e.g. flight simulation or Gatekeeper task) sometimes show increase in HRV [8,17] while tasks involving physical activity (e.g. biceps curls) can show decreases in HRV [71].

When it comes to the temporal factor for SBP and DBP, studies that have demonstrated sensitivity to fatigue have sometimes relied on repeated spot measurements (i.e., not continuous segments) [68] or compared changes in blood pressure before and after the fatigue task using repeated measurements rather than continuous segments [69]. However, to our knowledge, few studies have examined continuous BP in terms of prolonged task induced fatigue. HR has been measured using segments ranging from two to 15 minutes, while HRV has frequently been measured using segments ranging from one to 15 minutes [17,69,71,73] during tasks lasting from 10 minutes to one hour and 48 minutes.

In sum, HR and HRV, both of which can be measured continuously and non-invasively, are promising variables for indicating fatigue, though their interpretation may vary based on physical or mental demands. BP, both SBP and DBP, should also be considered. However, the literature on blood pressure is generally less comprehensive than that on HR, HRV, and fatigue, and sometimes relies on spot measures rather than continuous ones.

### The present study

There is a strong indication that cardiovascular variables, in particular HR and HRV, are sensitive to variations in mental states such as workload, fatigue, and stress. However, the sensitivity of these measures appears to depend on the task type, task duration, and the overall level of task demand and difficulty. It is therefore important to compare the sensitivity of HR, HRV and BP across a broader range of task types, cognitive demands, and time frames.

The aim of the present study is to extend the literature by measuring real-time cardiovascular reactivity across three mental states: cognitive workload, stress, and fatigue, by using several different tasks over a period of approximately 1 ½ to 2 hours. The cardiovascular variables measured include HR and BP (SBP and DBP) as well as HRV measures (IBI, RMSSD, and HF). Our cognitive workload condition will consist of four cognitive tasks that tap into visuospatial working memory, decision-making, and executive function. The stress condition will be a modified version of the TSST, consisting of three stress periods: preparation for speech, speech condition, and an arithmetic task. In the fatigue condition, we will use time on task for a mental task (as opposed to a physical task) where participants perform continuously on the Stroop task for 15 minutes and the Psychomotor vigilance task (PVT) for 10 minutes.

Three linear mixed models will be conducted for each cardiovascular variable: one for cognitive workload, one for stress, and one for fatigue. Based on the literature, we expect each cardiovascular variable to be sensitive to task-related differences, thereby differentiating at least one task from the reference task within each model. In addition, we will report each variable’s ability to differentiate between tasks as an indicator of sensitivity. Pairwise comparison of the estimated marginal means of the models will be reported using t-ratios and p-values corrected for multiple comparisons using Tukey’s method.

## Materials and methods

### Participants

The protocol of the study from which the data were obtained, including the recruitment process, cognitive tasks, equipment, and procedures, has been described in a prior publication [74]. Sixty individuals, 27 men (45%) and 33 women (55%), aged between 19 and 52 years old (*M* = 28.1 years, *SD* = 7.9), participated in the current study. Participants were recruited via online advertisements on Facebook and posters distributed in places such as gyms, swimming pools, and other public areas in Reykjavik. Furthermore, an email advertising the study was sent to all students at Reykjavik University.

The Icelandic National Bioethics Committee (VSN-21-087) approved the study. All participants read and signed an informed consent form before participation. Participants were encouraged to inform the researcher if they experienced any discomfort. Furthermore, participants were informed that they did not have to partake in individual assignments and could also discontinue their participation at any time. Although the risk of participating in the current study was very low, participants were informed that if they experienced any discomfort after participating, they could see a trained psychologist to discuss it. In return for participating, participants received a 4,000 ISK gift card for the local shopping center Kringlan.

### Materials

#### Workload: Visuospatial working memory

##### Judgment of Line Angle and Position (JLAP)

is a cognitive test that measures visuospatial capacity [75]. The task is adapted from Collaer [76], in which a target line is presented above a reference array. For 20 trials, each lasting 10 seconds, the participant’s goal is to match a target line to a reference array. The highest possible score for the total task is 20, corresponding to the number of trials [75].

##### The Corsi Block-Tapping Test

is a short-term memory test resembling a digit span test. The test’s name references its creator, Philip Michael Corsi, who developed it as a doctoral student [77]. The Corsi test assesses a person’s short-term vision and spatial memory. Originally, the test consisted of nine square blocks which were positioned on a wooden table. Since then, digital versions of the test have been created [78]. In the digital version used in the present study, the participant watches square blocks light up one after another in a specific order on a computer screen. When the sequence is over, the participant’s goal is to click on the boxes in the same order as they lit up.

#### Workload: Decision making

##### The Balloon Analogue Risk Task (BART)

The BART assesses decision making under uncertainty. Participants press a button to inflate a virtual balloon [79]. With each press, the balloon inflates further, and participants simultaneously see an increase in the simulated monetary award they have earned. However, each press can also cause the balloon to explode; if that happens, the monetary award is lost. For each pump, participants must decide whether to inflate the balloon further to earn more money, while risking making it explode and losing the money, or to cash out the money they have already made. Participants do not know when the balloon will explode and therefore, uncertainty always accompanies the decision. The BART requires a balance between the desire for a higher reward and the risk of loss under uncertainty. As in real world situations, participants can collect the monetary reward or leave the risky situation at any time, imitating decision making under real world risky circumstances. Therefore, BART has high validity [80,81].

#### Workload: Executive function - Inhibition and attentional control

##### The Stroop test

was developed by John Ridley Stroop and is a well-known instrument for neuropsychological measurement [82]. Although relatively simple to administer, it correlates significantly with other, more complex psychological measures [83]. Prior studies have used the Stroop task to assess different mental capacities, such as selective attention and cognitive control of automatic processes [20]. The Stroop test is divided into two scenarios. In the former, a colored word appears in ink that is incongruent to the word’s meaning (e.g., the word “blue” appears in “red” ink). In the latter scenario, the ink is congruent with the word’s meaning (e.g., the word “red” in “red” ink). The participant’s goal is to identify the ink color while ignoring the word’s meaning. The current study used three levels of the Stroop task: Time-limited (each word appeared for only a short period), partly incongruent (30% and 70% of the words matched the color), and 100% congruent (the color matched the word) [84]. The Stroop task in the workload condition lasted for five minutes, with the first minute used for analysis.

#### Stress

##### The Trier Social Stress Test (TSST)

was developed to study stress in a laboratory setting [85]. In a modified version of the test used in the current study, participants are told they have received an imaginary invitation to interview for their dream summer job. They are told that their speech will be recorded on camera and later watched and assessed by the researcher and three members of a hiring committee. The participant’s role is to convince the viewers that they are the most qualified person for the job. The participant gets three minutes to prepare their speech. While preparing, they are allowed to jot down speaking points; however, during the speech itself, they are forbidden to look at them. The participant is then informed that the recording will last four minutes and is asked to deliver the speech for the entire duration. If the participant finishes their speech before the time has passed, they are asked to continue speaking. Once the time has passed, the participant is told that their mathematical skills must be assessed for the recruitment. They are then given three minutes to subtract 17 from 2023 as often as they can, while the researcher listens and evaluates their performance. If they make a mistake during the subtraction, they are stopped and instructed to start over from 2023.

#### Fatigue

##### Stroop Fatigue (SF)

The Stroop test, described above, has previously been used to measure fatigue [86]. The current study used SF to measure fatigue over time. Participants performed the task (described above) for 15 minutes. The duration of the recordings varied due to noise, resulting in incomplete recordings for some participants. Therefore, the first 12 minutes were used since most participants had data available for this interval.

##### The Psychomotor Vigilance Task (PVT)

is a cognitive task originally created to assess sustained attention [87]. It has since become a widely used neuropsychological task [88,89]. During the 10-minute task, the participant sits in front of a computer screen and tries to respond as quickly as possible by pressing a button whenever a red square appears. These multiple trials provide a measure of sustained attention and reaction time [87]. In addition, PVT has been shown to be a reliable assessment of vigilance and attention [90]. Similar to the Stroop task, the PVT has also been used to assess mental fatigue through time-on-task-related vigilance decrements (e.g., slowed reaction times) over prolonged performance [87,89,90].

### Equipment

While participants underwent the cognitive, stress, and fatigue tasks, HR, BP, and IBI were continuously recorded using a CareTaker medical device [91]. The CareTaker is a noninvasive tool that measures vital signs such as HR, BP, IBI, and respiratory rate. The CareTaker measures the beat-to-beat arterial pulse-pressure waveform using pulse decomposition analysis through a low-pressure inflatable cuff placed on the middle fingertip [92]. The device has been more extensively described in prior studies [92,93] and has been cleared by the FDA for continuous monitoring of HR, BP, and respiratory rate in adults (FDA K163255). In the current study, to calibrate the CareTaker, participants’ blood pressure was measured using a cuff placed on the upper arm. The CareTaker device was then placed on the participant’s wrist, and the cuff was placed on the middle finger. The SBP and DBP values obtained from upper-arm blood pressure measurements were then entered into an electronic tablet to calibrate the CareTaker. The same tablet was used to record timestamps for the tasks using the CareTaker app. Prior studies have validated the CareTaker by correlating its measures with invasive arterial pressure measurements. For instance, Kwon et al. [92] found that the CareTaker accurately tracked beat-to-beat BP, and that the CareTaker IBI was highly correlated with the IBI obtained from the invasive arterial pressure waveform.

### Procedure

The current study used a subset of data from a larger study conducted in 2021. Participants attended a laboratory at Reykjavik University. After providing informed consent, participants had their blood pressure measured. After measuring blood pressure, the researcher calibrated the CareTaker to record real-time physiological data during the study. Participants were seated in front of a computer screen and asked to keep their hand as still as possible to ensure the quality of the data recorded by the CareTaker. Cardiovascular reactivity was measured using a small inflatable cuff placed on the participant’s finger. First, a three-minute baseline was recorded for each participant. After the baseline period, the participants underwent various cognitive and neuropsychological tests, not all of which were included in the current study. The cognitive (workload) tests used in the current study were performed in the following order: JLAP, CORSI, BART, and Stroop. The time required to complete the tests varied between participants. The cognitive tests took at least 25 minutes to complete, the TSST around ten minutes, the SF around 15 minutes, and the PVT ten minutes. However, when instructions, clarifications, equipment adjustments, and transitions between tasks are taken into account, the experiment lasted about one and a half to two hours.

### Extraction of cardiovascular measures

The continuous cardiovascular data were extracted from the Caretaker device using Microsoft Excel. Each task, including the baseline, was identified manually by matching the task timestamps with the Caretaker output.

Averages for HR, SBP, DBP, and IBI were then calculated for each task and subtracted from the corresponding baseline measure, creating a difference score for each variable. For JLAP, Corsi, BART, Stroop and TSST, the first minute for each task was used to capture task-specific changes in cardiovascular reactivity, immediately after task onset. In contrast, the aim for the fatigue condition was to examine changes in cardiovascular reactivity over time. Therefore, both the SF and PVT tasks were divided into three equal intervals (approximately three minutes for PVT and four minutes for SF). The values were then subtracted from baseline (difference scores).

For HRV analysis, IBI data for each task were exported to .txt format and analyzed using Kubios Scientific (version 4.1.2.1). Current guidelines recommend a recording interval of five minutes for reliable estimation of RMSSD and at least one minute for HF measures [49]. Although longer segments are preferable, one-minute recordings have been shown to be efficient for RMSSD, i.e., correlating strongly with longer recordings [94–96]. Still, given the well-established five-minute recommendation for short-term RMSSD recordings [48,49], the current study used the five-minute intervals where possible. However, since only Stroop, BART, and, in some instances, Corsi reached a five-minute interval, the entire task duration was used for the other tasks that did not reach the five-minute threshold. The durations of the cognitive tasks ranged from one minute and 37 seconds to five minutes, and segments shorter than one minute were excluded. For the fatigue condition, PVT (10 min) and SF (12 min) were divided into equal intervals to indicate fatigue over time. After all HRV recordings had been imported into Kubios, each recording was visually inspected for artifacts, and thereafter, the default automatic artifact correction in Kubios was applied. Recordings with more than 10% corrected beats were excluded, matching an approach used in prior studies [97,98]. The automatic artifact correction in Kubios corrects both missed and ectopic beats using a cubic-spline interpolation method [99]. As with HR, IBI, SBP, and DBP, difference scores for the HRV variables were calculated in Microsoft Excel by subtracting baseline values from task values.

### Data analysis

For the statistical analysis, the current study used linear mixed models (LMMs), an approach widely used in psychophysiology for repeated cardiovascular measures nested within participants [100,101]. One reason for using this approach is that LMMs can handle missing data better than typical repeated measures ANOVA or MANOVA [102]. For instance, the model keeps participants in the analysis even when they have missing values, thereby maximizing the use of available data. In addition, individual variability can be accounted for by including random effects, which can reduce noise from individual differences (e.g., between low and high HR). Since our dataset had missing values due to noise, resulting in some participants having missing data, as well as measurements of cardiovascular reactivity, which are known to have substantial individual variability, we chose LMMs to account for that.

Three LMMs were conducted for each cardiovascular variable: one for the cognitive condition (workload), one for the stress condition (TSST), and one for the fatigue condition. Tasks and gender (1 = male, 2 = female) were coded as factor variables. Each model was built using the following structure lmer(cardiovascular variable ∼ task + age (centered) + gender + (1 | Subject), data = data). Random intercepts were included for each participant to account for individual variability in the cardiovascular measures. In addition, age (mean-centered) and gender were added as covariates. Finally, estimated marginal means (emmeans) and Tukey post hoc correction for multiple pairwise comparisons were used to report differences between individual pairs of tasks; the significance threshold was set to *p* < .05. All data were analyzed using R (version 4.4.1). The packages used for the analysis were tidyverse, lme4, emmeans, ggplot2, patchwork, and dplyr. Due to missing data, the number of participants differed across some models. Therefore, to provide a cleaner overview of the analysis, descriptive statistics such as the number of participants, mean age, and gender ratio are reported for each model. Similar to Schwerdtfeger and Schlagert [103], we used t > 2 as a heuristic for statistical significance in the LMMs, due to known challenges in estimating *p* values for fixed effects. However, since p-values are standard in statistical reporting, we also report p-values estimated from marginal means for pairwise comparisons. The ability of each cardiovascular measure to differentiate between tasks was interpreted as sensitivity.

## Results

We used LMM to examine the effect of different tasks on cardiovascular reactivity. Separate analyses were conducted for the cognitive workload condition (JLAP, CORSI, BART, and Stroop), the stress condition (TSST [Preparation, Speech, Math]), and the fatigue condition (Stroop and PVT, each divided into three equal parts). Due to variations in missing data, the models include different subsets of participants. Therefore, we report the number of participants (*n*), mean age (*M*_age_), standard deviation (SD), and gender ratio for each model. Reference tasks were selected in alphabetical order in R. Since males were used as the reference group for gender, each intercept represents the estimated value for male participants. Since our analyses rely on difference scores and the main focus is on pairwise comparisons of estimated marginal means, the reference task is not of particular interpretive importance. However, we do report significant results from the models for transparency. Reporting will follow this order: for each cardiovascular variable, unadjusted coefficients from the LMMs will be reported in the text, along with t-values. Next, results from the estimated marginal means for all relevant conditions (cognitive workload, stress, and fatigue) will be presented in a figure and a table, with Tukey corrections (*p* values) for multiple comparisons. Since age (centered) and gender did not yield significant effects in any model, statistics for these variables will not be reported in the results section to streamline reporting. However, full regression model results (including *b*, *SE,* 95% CI*, df, t,* and *p)* and pairwise comparisons (including mean difference, 95% CI*, SE df, t,* and *p*) for all conditions and variables are available in the Supplementary Material (Tables S1-S18).

### Models for heart rate

For the cognitive tasks, the LMM (before correcting for multiple comparisons) for HR (*n* = 42, *M*_age_ = 28, *SD* = 7.70, 52.4% female) showed significantly increased HR during JLAP (*b* = 3.84, *SE* = 0.87, *t* = 4.42), Corsi (*b* = 3.90, *SE* = 0.88, *t* = 4.46) and Stroop (*b* = 3.66, *SE* = 0.98, *t* = 3.74) compared to the reference task BART.

For the TSST, the LMM (before correcting for multiple comparisons) for HR (*n* = 37, *M*_age_ = 28.22, *SD* = 7.79, 54.1% female) showed that HR increased significantly (*t* > 2) during Preparation (*b* = 3.06, *SE* = 1.19, *t* = 2.58) and Speech condition (*b* = 6.30, *SE* = 1.21, *t* = 5.22) compared to the reference task Math.

The LMM for HR (before correcting for multiple comparisons) during the fatigue condition (*n* = 53, *M*_age_ = 28.40, *SD* = *7.94,* 54.72% female) showed that SF part one (*b* = 6.12, *SE* = 0.97, *t* = 6.31), SF part two (*b* = 6.02, *SE* = 0.97, *t* = 6.20) and SF part three (*b* = 5.65, *SE* = 0.97, *t* = 5.82) were all significantly increased (*t* > 2) compared to the reference group, PVT part one. However, PVT parts two and three did not differ significantly from PVT part one. Fig 1 shows the estimated marginal means of HR for each task condition, derived from the LMM using emmeans, adjusted for gender and age.

**Fig 1.**
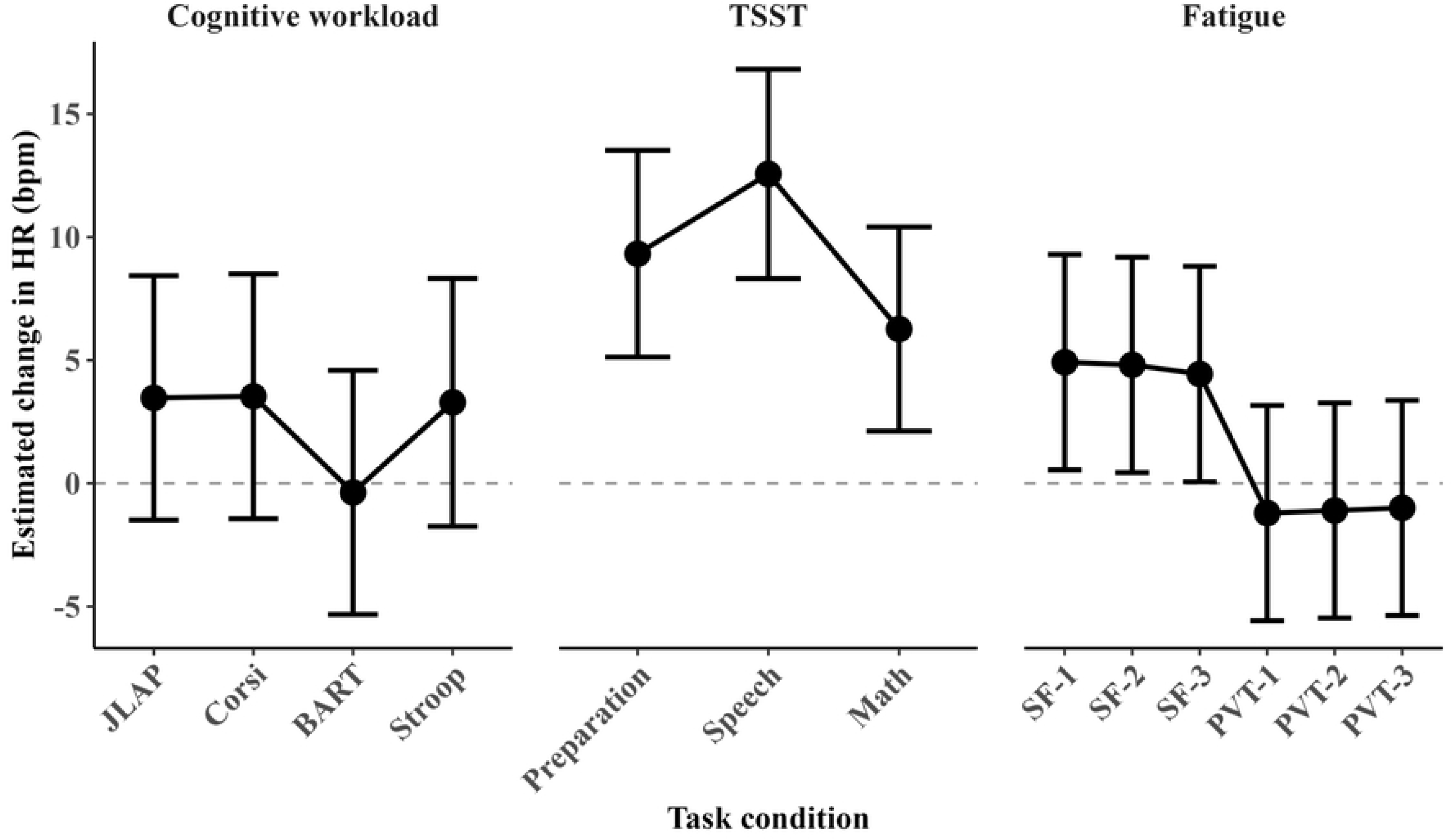
Estimated marginal means for HR from the three models adjusted for age and gender, with 95% confidence intervals. *Note:* The estimates were obtained using estimated marginal means (emmeans); the dashed line indicates no change from baseline. bpm = beats per minute, JLAP = judgment of line angle and position; Corsi = Corsi block-tapping task; BART = balloon analogue risk task; Stroop = Stroop task; TSST = Trier social stress test; SF = Stroop fatigue intervals one, two and three; PVT = psychomotor vigilance task intervals one, two, and three. Numbers indicate the three equal parts of each task in chronological order.

To examine HR’s ability to differentiate between tasks, pairwise comparisons were conducted using estimated marginal means (emmeans) from each model, adjusted for age and gender. Tukey was used to correct for multiple comparisons. Significant pairwise comparisons from the models are presented in Table 1, with Tukey adjustment for multiple comparisons.

**Table 1.** Pairwise comparisons (Tukey-adjusted) from the three HR models adjusted for age and gender.

| Model | *n* | Comparison | Estimated mean difference (bpm) | 95% CI | *SE* | *df* | *t* ratio | *p* |
| --- | --- | --- | --- | --- | --- | --- | --- | --- |
| Cognitive workload | 42 | BART – Corsi | -3.90 | [-6.19, -1.61] | 0.88 | 99.17 | -4.46 | < .001*** |
| Cognitive workload | 42 | BART – JLAP | -3.84 | [-6.11, -1.57] | 0.87 | 99.30 | -4.42 | < .001*** |
| Cognitive workload | 42 | BART – Stroop | -3.66 | [-6.22, -1.10] | 0.98 | 99.62 | -3.73 | .002** |
| TSST | 37 | Math – Preparation | -3.06 | [-5.93, -0.19] | 1.19 | 50.45 | -2.57 | .034* |
| TSST | 37 | Math – Speech | -6.30 | [-9.22, -3.38] | 1.21 | 50.02 | -5.21 | < .001*** |
| TSST | 37 | Preparation – Speech | -3.25 | [-6.37, -0.12] | 1.29 | 50.77 | -2.51 | .040* |
| Fatigue | 53 | PVT-1 - SF-1 | -6.12 | [-8.91, -3.34] | 0.97 | 257.05 | -6.31 | < .001*** |
| Fatigue | 53 | PVT-1 - SF-2 | -6.02 | [-8.81, -3.23] | 0.97 | 257.05 | -6.20 | < .001*** |
| Fatigue | 53 | PVT-1 - SF-3 | -5.65 | [-8.44, -2.86] | 0.97 | 257.05 | -5.82 | < .001*** |
| Fatigue | 53 | PVT-2 - SF-1 | -6.02 | [-8.81, -3.23] | 0.97 | 257.05 | -6.20 | < .001*** |
| Fatigue | 53 | PVT-2 - SF-2 | -5.92 | [-8.71, -3.13] | 0.97 | 257.05 | -6.09 | < .001*** |
| Fatigue | 53 | PVT-2 - SF-3 | -5.55 | [-8.33, -2.76] | 0.97 | 257.05 | -5.71 | < .001*** |
| Fatigue | 53 | PVT-3 - SF-1 | -5.91 | [-8.70, -3.13] | 0.97 | 257.05 | -6.09 | < .001*** |
| Fatigue | 53 | PVT-3 - SF-2 | -5.81 | [-8.60, -3.02] | 0.97 | 257.05 | -5.98 | < .001*** |
| Fatigue | 53 | PVT-3 - SF-3 | -5.44 | [-8.23, -2.65] | 0.97 | 257.05 | -5.60 | < .001*** |

*Note: n* = number of participants for each model; bpm = beats per minute; 95% CI = 95%
confidence intervals; *df* = degrees of freedom; *SE* = *standard error*; 95% confidence intervals
and *p* values were adjusted using the Tukey method. Estimated mean differences and 95%
confidence intervals are presented as bpm. \**p* < .05; \*\**p* < .01; \*\*\**p* < .001.

### Models for interbeat interval

The IBI LMM for the cognitive tasks (*n* = 42, *M*_age_ = 28, *SD* = 7.70, 52.40% female), showed significant decrease in IBI (ms) during JLAP (*b* = -32.55, *SE* = 9.47, *t* = -3.44), Corsi (*b* = -42.37, *SE* = 9.56, t = -4.43) and Stroop (*b* = -45.04, *SE* = 10.64, *t* = -4.23) compared with the reference task BART.

The IBI LMM for TSST (*n* = 37, *M*_age_ = 28.22, *SD =* 7.79, 54.1% female) showed a significant decrease in IBI between the speech condition (*b* = -34.08, *SE* = 7.45, *t* = -4.57) and the reference task, math. However, there was no significant difference between the preparation condition (b = -4.33, SE = 7.32, t = -0.59) and the reference task.

The IBI LMM for the fatigue condition (*n* = 49, *M*_age_ = 28.12, *SD* = 7.55, female = 53.06%) showed that SF part one (*b* = -53.14, *SE* = 8.36, *t* = -6.35), SF part two (*b* = -52.47, *SE* = 8.36, *t* = -6.27) and SF part three (*b* = -50.12, *SE* = 8.36, *t* = -5.99) showed significant decrease in IBI (ms) from the reference task PVT part one. However, PVT parts two and three did not differ from the reference task, PVT part one. Fig 2 shows estimated marginal means (emmeans) of IBI for each model by task condition, derived from the LMM, adjusted for gender and age.

**Fig 2.**
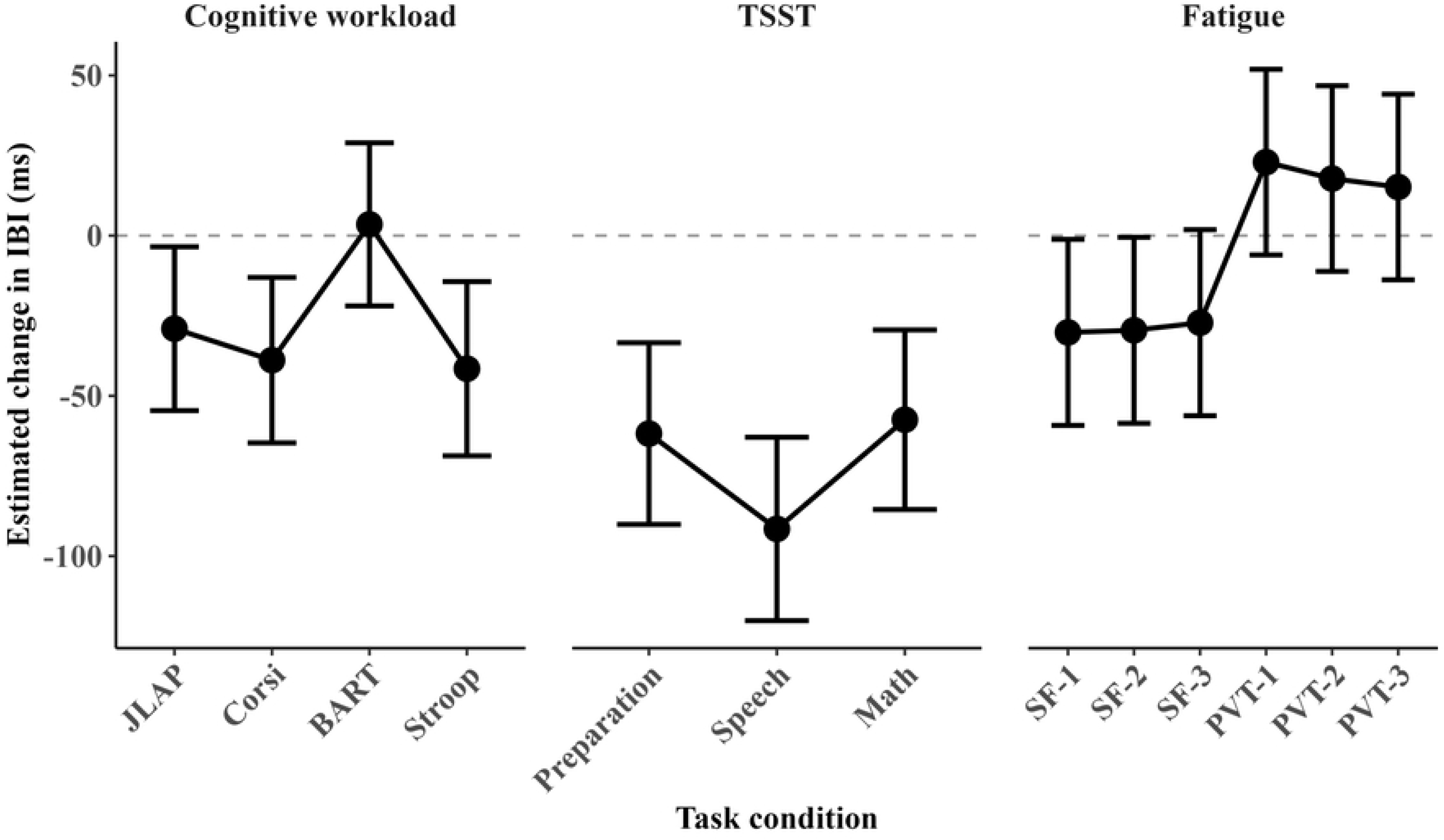
Estimated marginal means for IBI from the three models adjusted for age and gender, with 95% confidence intervals. *Note:* The estimates were obtained using estimated marginal means (emmeans); the dashed line indicates no change from baseline. ms = milliseconds, JLAP = judgment of line angle and position; Corsi = Corsi block-tapping task; BART = balloon analogue risk task; Stroop = Stroop task; TSST = Trier social stress test; SF = Stroop fatigue intervals one, two, and three; PVT = psychomotor vigilance task intervals one, two, and three. Numbers indicate the three equal parts of each task in chronological order.

Similar to HR, we also used pairwise comparisons on estimated marginal means to examine the ability of IBI to differentiate between tasks. Pairwise comparisons were conducted using estimated marginal means (emmeans) from each model, adjusting for age and gender. Tukey correction was used to correct for multiple comparisons. Significant pairwise comparisons from the models are presented in Table 2, with Tukey adjustment for multiple comparisons.

**Table 2.** Pairwise Comparisons for Each Model Corrected with Tukey.

| Model | *n* | Comparison | Estimated mean difference (ms) | 95% CI | *SE* | *df* | *t* ratio | *p* |
| --- | --- | --- | --- | --- | --- | --- | --- | --- |
| Cognitive workload | 42 | BART - Corsi | 42.37 | [17.39, 67.35] | 9.56 | 99.79 | 4.43 | < .001*** |
| Cognitive workload | 42 | BART - JLAP | 32.55 | [7.81, 57.29] | 9.47 | 100.29 | 3.44 | .005** |
| Cognitive workload | 42 | BART - Stroop | 45.04 | [17.22, 72.86] | 10.65 | 101.81 | 4.23 | < .001*** |
| TSST | 37 | Math - Speech | 34.08 | [16.07, 52.09] | 7.46 | 49.84 | 4.57 | < .001*** |
| TSST | 37 | Preparation - Speech | 29.74 | [10.46, 49.03] | 7.99 | 50.47 | 3.72 | .001** |
| Fatigue | 49 | PVT-1 - SF-1 | 53.14 | [29.10, 77.17] | 8.36 | 237.10 | 6.35 | < .001*** |
| Fatigue | 49 | PVT-1 - SF-2 | 52.47 | [28.44, 76.50] | 8.36 | 237.10 | 6.27 | < .001*** |
| Fatigue | 49 | PVT-1 - SF-3 | 50.12 | [26.09, 74.15] | 8.36 | 237.10 | 5.99 | < .001*** |
| Fatigue | 49 | PVT-2 - SF-1 | 48.01 | [23.98, 72.05] | 8.36 | 237.10 | 5.74 | < .001*** |
| Fatigue | 49 | PVT-2 - SF-2 | 47.35 | [23.31, 71.38] | 8.36 | 237.10 | 5.66 | < .001*** |
| Fatigue | 49 | PVT-2 - SF-3 | 45.00 | [20.97, 69.03] | 8.36 | 237.10 | 5.38 | < .001*** |
| Fatigue | 49 | PVT-3 - SF-1 | 45.38 | [21.35, 69.41] | 8.36 | 237.10 | 5.43 | < .001*** |
| Fatigue | 49 | PVT-3 - SF-2 | 44.71 | [20.68, 68.74] | 8.36 | 237.10 | 5.35 | < .001*** |
| Fatigue | 49 | PVT-3 - SF-3 | 42.36 | [18.33, 66.40] | 8.36 | 237.10 | 5.06 | < .001*** |

*Note:* *n* = number of participants for each model; ms = milliseconds; 95% CI = 95% confidence intervals; *df* = degrees of freedom; *SE* = standard error; 95% confidence intervals and *p* values were adjusted using the Tukey method. Estimated mean differences and 95% confidence intervals are presented as ms. \**p* < .05; \*\**p* < .01; \*\*\**p* < .001.

### Models for systolic and diastolic blood pressure

Similar to HR and IBI, six separate LMMs were run for SBP and DBP (covering cognitive, stress, and fatigue conditions). None of these models yielded significant results after correcting for multiple comparisons and are therefore not presented in detail. However, results showed that SBP was significant in the preparation condition of TSST (*b* = -1.65, *SE* = 0.80, *t* = - 2.06), compared to the reference task TSST math, but only before, but not after correction for multiple comparisons (*p* = .110). Similar results were found for the same tasks in DBP as well (b = -1.69, SE = 0.80, t = -2.11), which were also not significant after correcting for multiple comparisons (p = .098).

### Models for HRV-RMSSD

The LMM for HRV-RMSSD (ms) for the cognitive tasks (*n* = 42, *M*_age_ = 28, *SD* = 7.70, female = 52.38%) showed that HRV-RMSSD was only able to differentiate between Stroop (*b* = 9.22, *SE* = 2.22, *t* = 4.16) and the reference task BART. However, after estimating the marginal means and correcting for multiple comparisons, Stroop also differed significantly from Corsi and JLAP (*p* < .001).

In the LMM for HRV-RMSSD during the TSST, none of the conditions (preparation, speech, or math) differed significantly. Therefore, the results from this model are not presented in more detail but, like all other models, are available in the Supplementary Material (Table S14).

The LMM for HRV-RMSSD during the fatigue conditions (*n* = 32, *M*_age_ = 30, *SD* = 8.51, female = 62.50%) showed that PVT part three (*b =* -8.33, *SE* = 2.95, *t* = -2.82), SF part one (*b =* -6.73, *SE* = 3.14, *t* = -2.14), SF part two (*b =* -9.08, *SE* = 3.14, *t* = -2.90) and SF part three (*b =* - 8.99, *SE* = 3.14, *t* = -2.87) significantly decreased compared to the reference group which was PVT part one. However, after correcting for multiple comparisons using emmeans and Tukey correction, none of these comparisons remained significant. Fig 3 shows the estimated marginal means of RMSSD and 95% confidence intervals for each task condition, derived from the LMM using emmeans and adjusted for age and gender

**Fig 3.**
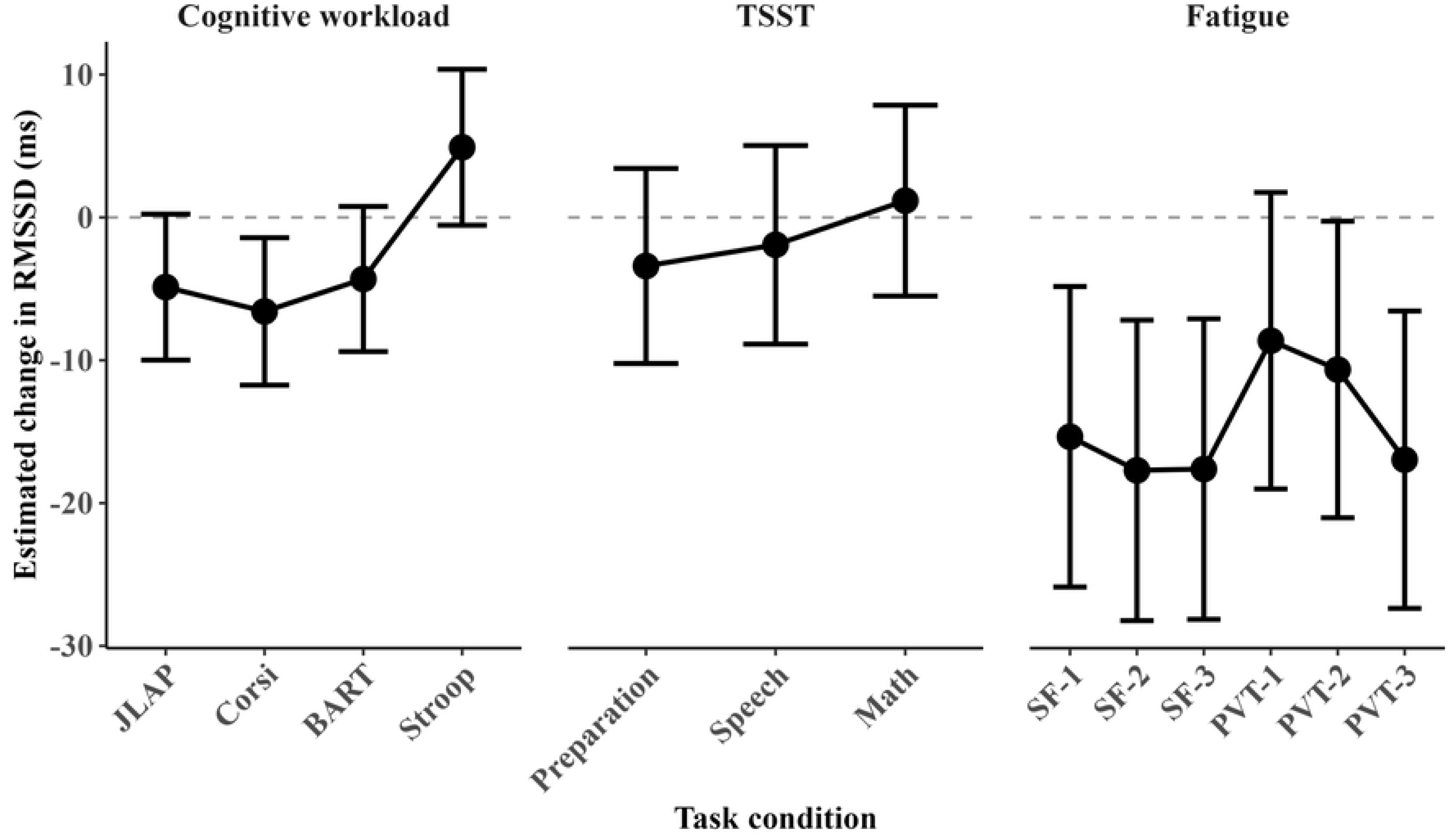
Estimated marginal means for RMSSD (ms) from the three models adjusted for age and gender, with 95% confidence intervals. *Note:* The estimates were obtained using estimated marginal means (emmeans); the dashed line indicates no change from baseline. ms = milliseconds, JLAP = judgment of line angle and position; Corsi = Corsi block-tapping task; BART = balloon analogue risk task; Stroop = Stroop task; TSST = Trier social stress test; SF = Stroop fatigue intervals one, two and three; PVT = psychomotor vigilance task intervals one, two, and three. Numbers indicate the three equal parts of each task in chronological order.

To assess how well RMSSD differentiated between the tasks, we conducted pairwise comparisons using estimated marginal means (emmeans) from each model. Significant pairwise comparisons from the models are presented in Table 3, with Tukey adjustment for multiple comparisons.

**Table 3.** Pairwise comparisons (Tukey-adjusted) from the three RMSSD models adjusted for age and gender.

| Model | *n* | Comparison | Estimated mean difference (ms) | 95% CI | *SE* | *df* | *t* ratio | *p* |
| --- | --- | --- | --- | --- | --- | --- | --- | --- |
| Cognitive workload | 42 | BART - Stroop | -9.22 | [-15.02, -3.43] | 2.22 | 102.07 | -4.15 | < .001*** |
| Cognitive workload | 42 | Corsi - Stroop | -11.49 | [-17.40, -5.59] | 2.26 | 101.96 | -5.08 | < .001*** |
| Cognitive workload | 42 | JLAP - Stroop | -9.79 | [-15.60, -3.97] | 2.23 | 101.93 | -4.40 | < .001*** |
| Fatigue | 32 | PVT-1 - PVT-3 | 8.33 | [-0.21, 16.87] | 2.95 | 124.04 | 2.82 | .060 |
| Fatigue | 32 | PVT-1 - SF-1 | 6.73 | [-2.35, 15.81] | 3.14 | 125.12 | 2.14 | .272 |
| Fatigue | 32 | PVT-1 - SF-2 | 9.08 | [-0.00, 18.16] | 3.14 | 125.12 | 2.89 | .050 |
| Fatigue | 32 | PVT-1 - SF-3 | 8.99 | [-0.09, 18.07] | 3.14 | 125.12 | 2.87 | .054 |
| Fatigue | 32 | PVT-2 - PVT-3 | 6.31 | [-2.22, 14.85] | 2.95 | 124.04 | 2.14 | .274 |
| Fatigue | 32 | PVT-2 - SF-2 | 7.06 | [-2.02, 16.14] | 3.14 | 125.12 | 2.25 | .222 |
| Fatigue | 32 | PVT-2 - SF-3 | 6.97 | [-2.11, 16.05] | 3.14 | 125.12 | 2.22 | .235 |

*Note:* *n* = number of participants for each model; ms = milliseconds; 95% CI = 95% confidence intervals; *df* = degrees of freedom; *SE* = standard error; 95% confidence intervals and *p* values were adjusted using the Tukey method. Estimated mean differences and 95% confidence intervals are presented as ms. \**p* < .05; \*\**p* < .01; \*\*\**p* < .001.

### Models for high frequency HRV

The LMM for the cognitive tasks (*n* = 42, *M*_age_ = 28, *SD* = 7.70, 52.38% female) showed (before correcting for multiple comparisons) that HRV-HF absolute power ms^2^ (0.15 – 0.4 Hz) was decreased (ms²) during the Corsi task (*b* = -221.40, *SE* = 107.31, *t* = -2.06) and increased during the Stroop task (*b* = 243, *SE* = 119.35, *t* = 2.04), compared with the reference task which was BART. After correcting for multiple comparisons, Stroop vs Corsi was significant (*p* = .001) as well as JLAP vs Stroop (*p* = .01).

As with the other cardiovascular variables, three LMMs were fitted to high-frequency power. The TSST and fatigue models did not yield significant results either in the model or in the pairwise comparison; therefore, they are not presented in detail in the current paper. However, as with all other regression models conducted for the current study, they are presented in the Supplementary Material (Tables S17-S18).

Fig 4 shows estimated marginal means for HRV-HF for all three models derived from the LMM using emmeans, adjusted for gender and age.

**Fig 4.**
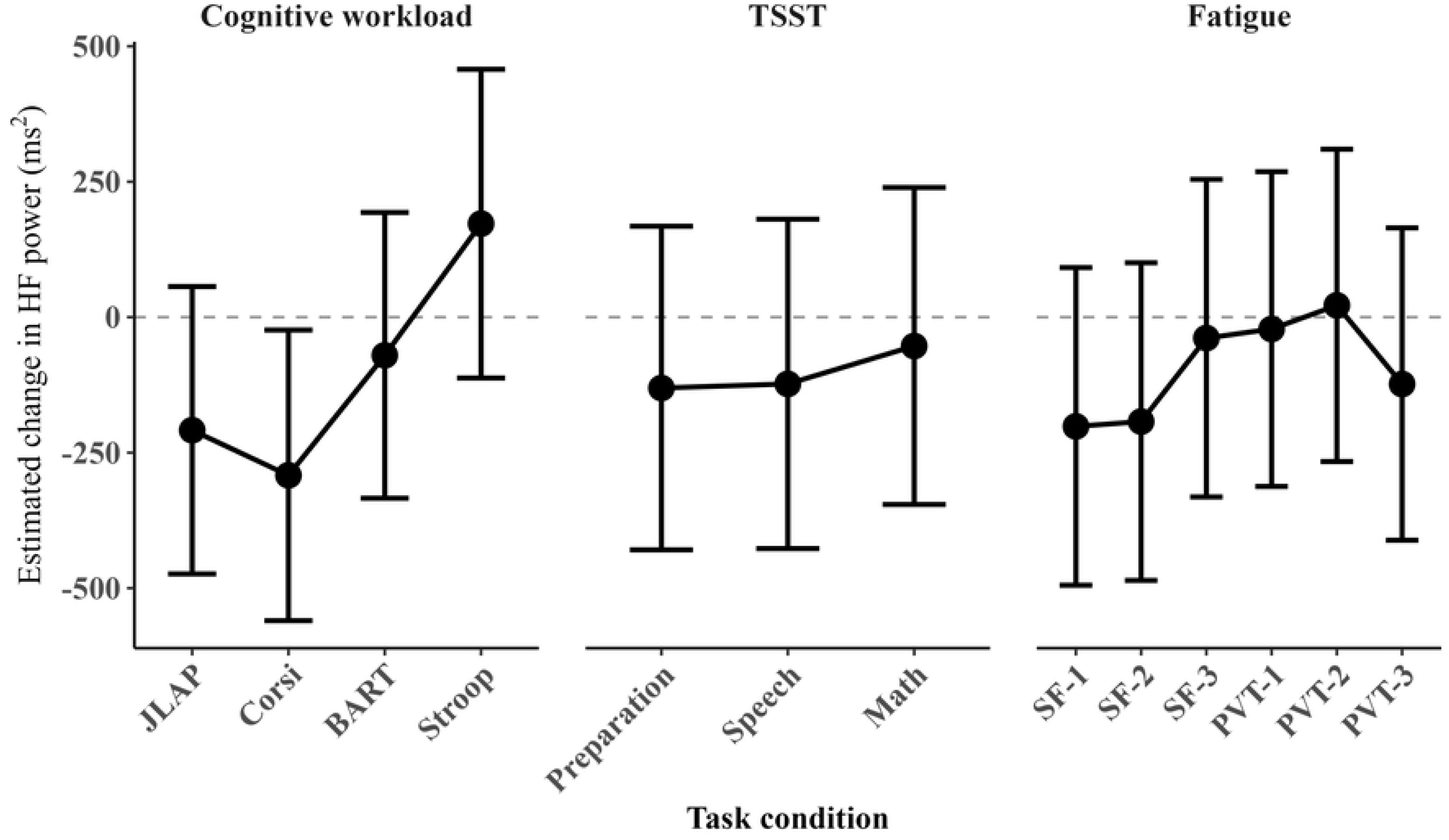
Estimated marginal means for HF absolute power ms^2^ from the three models adjusted for age and gender, with 95% confidence intervals. *Note:* The estimates were obtained using estimated marginal means (emmeans); the dashed line indicates no change from baseline. ms^2^ = milliseconds squared, JLAP = judgment of line angle and position; Corsi = Corsi block-tapping task; BART = balloon analogue risk task; Stroop = Stroop task; TSST = Trier social stress test; SF = Stroop fatigue intervals one, two and three; PVT = psychomotor vigilance task intervals one, two, and three. Numbers indicate the three equal parts of each task in chronological order.

## Discussion

The current study aimed to identify which cardiovascular variables were the most sensitive to detecting different mental states (cognitive workload, stress, and fatigue). We used three separate linear mixed models for each cardiovascular variable. Sensitivity was defined as the ability of the cardiovascular variable to differentiate between the conditions. The ability to differentiate between the tasks was detected using *t* values from the models and *p* values from pairwise comparisons of the estimated marginal means from the models, corrected for multiple comparisons with Tukey’s method. Four different cognitive tasks (JLAP, Corsi, BART, and Stroop) were used to measure cognitive workload, TSST was used to measure stress, and the fatigue condition consisted of Stroop and PVT, each divided into three equal parts to capture fatigue over time. Instead of creating multiple hypotheses, we used general expectations that each cardiovascular variable would be able to differentiate between the task conditions.

### HR

In accordance with previous literature, we expected HR to be sensitive to differentiating between cognitive tasks. In addition, we expected HR to differentiate between all three TSST conditions and be sensitive to detect fatigue over time during the SF and PVT tasks.

The results for HR showed significant differences between conditions and their reference task in all three models (cognitive, TSST, and fatigue). Corrected pairwise comparisons showed that HR was sensitive in differentiating between some, but not all, tasks in each model. HR has previously been used successfully as a sensitive measure of different mental states, such as increased cognitive workload, stress, and fatigue [8,15,20,28]. HR may therefore be a good choice when identifying sensitive measures of different mental states. In the current study, HR significantly differentiated between all conditions in the TSST. These results align with previous findings on robust stress. Other studies have also found that HR distinguishes well between the different stages of TSST [15]. HR has also been found to detect robust stress in situations such as a simulation of complex military operations [55,56].

HR may be an overall good metric for detecting both robust and less robust stressors [15]. However, this may be dependent on the task, a recent study, using machine learning to classify cognitive workload and stress, found that adding distracting sounds to an n-back was not enough to classify stress using HR [14].

Although HR was clearly sensitive to stress in the current study, HR was not able to differentiate between all the cognitive workload tasks in the pairwise comparisons. HR could differentiate between BART and the other cognitive tests, but other comparisons were insignificant. Interestingly, the difference score was lowest during BART, indicating that HR was closest to baseline compared with the other tasks. A possible reason for this could be that BART is a relatively passive test, relying more on strategy and long-term memory compared with the other tests (JLAP, Corsi, and Stroop), which rely on constant attention, inhibition, and working memory. Prior work has suggested that HR may be more responsive to tasks relying on sustained attention and working memory [21,39]. Therefore, HR may be sensitive to distinguishing dynamic changes in workload that depend more on attentional and working memory processes rather than workload linked with decision-making and problem-solving tasks, where changes in workload are slow and perhaps cumulative over time. In fact, prior work has shown that HR does not well detect slow-paced, more problem-solving based tasks from baseline [3].

In the current study, fatigue was induced using a 15-minute Stroop task (SF) and an additional 10-minute PVT task at the end of the entire task setup (one and a half to two hours in total). The results showed an overall increase in HR across all three SF time points compared to the three PVT time points, and significant differences between each SF and each PVT time point. A systematic review of 30 studies examining fatigue detection during simulated or real-world driving found that fatigue was associated with decreased HR during prolonged task performance (typically lasting between 60 and 120 minutes) [13]. To our surprise, relatively few studies seem to directly address the role of HR on mental fatigue with time-on-task effect. However, at least one study showed a clear and gradual decrease in HR using cognitive tasks such as bimodal 2-back that lasted for around 2 hours with a 12-minute break [17]. These results align with the results of the current study, which indicate increased fatigue (decreased HR) at the end of the experiment, as the PVT was the last task for participants. A study by Mascord and Heath [104] used a visual tracking task to induce fatigue, combined with a reaction-time task, over a period of 140 minutes. Their results showed that HR began to decrease significantly after minute 105 and decreased by 8.7 bpm from the start of the experiment until the end. However, this pattern is not replicated in all studies. For instance, Karthikeyan et al. [105] found HR to increase over a prolonged 60-minute visuospatial working memory (2-back) task, suggesting that task type plays a role in cardiovascular response to fatigue. Therefore, it should also be considered that the differences between task types (SF and PVT) may have contributed to the effect. The study by Karthikeyan et al. [105] measured effort subjectively and found that effort did not reduce over the course of the experiment, indicating that participants put effort into staying alert, while fatigue, in a way, reflects surrendering. Our results may reflect a difference between more effortful cognitively demanding Stroop and more monotonous PVT. Future work on fatigue detection needs to take both type of fatigue inducing task and duration into consideration.

### HRV-IBI

As the fundamental beat-to-beat time-domain signal underlying HRV computation, the interbeat interval (IBI) provides a direct measure of cardiac timing and forms the basis for derived HRV indices such as RMSSD and HF. In the present study, we expected IBI to be sensitive to differences across the cognitive workload, stress, and fatigue conditions. Similar to HR, IBI differentiated between the reference task and several experimental conditions across all three models. The pairwise comparison was very similar to the HR comparison; that is, IBI could differentiate between BART and the other cognitive tasks, but the other tasks (JLAP, Corsi, and Stroop), did not differ significantly in their IBI profile. These results align with previous studies, which have found that IBI is a sensitive measure of cognitive workload, but its sensitivity seems to vary across studies [10,18] and may depend on the task type, similar to HR. For example, in a study in which participants used a driving simulator to perform n-back tasks at three levels, the aim was to mimic different levels of multitasking. The results showed that IBI was more sensitive than HRV-RMSSD, which could differentiate between two conditions, while IBI could differentiate between all three [18]. The same study found that other HRV measurements were altered by increased cognitive workload but were insignificant.

Another study that used IBI as a measure of increased cognitive workload among 27 pilots in the Finnish Air Force showed that IBI could differentiate between baseline and three performance conditions: low, medium, and high [10]. The same study could differentiate between high and medium, but not between low and medium conditions. It could be argued that these results are partly similar to the current study, since the current study found statistical differences between many of the cognitive tests but not all. This indicates that although IBI is a sensitive measure for increased workload, like with HR and other cardiovascular variables, it should be used with caution to indicate cognitive workload, especially when used by itself.

For the TSST, IBI (*p* = .001) outperformed HR (*p* = .040) in terms of the ability to differentiate between Preparation and Speech and performed equally when it came to differentiating between Math and Speech (*p* < .001). However, only HR was able to differentiate between Math and Preparation (*p* = .034), whereas IBI did not (*p* = .825). This indicates that even though IBI is an inverse (decreases as HR increases) of HR, both variables might contribute different information between more subtle changes in stress. This emphasizes our point that although some variables are sensitive, their sensitivity can vary by situation, and they should be used with caution, preferably not alone.

The fatigue condition also yielded results similar to HR; that is, IBI was higher during all three PVT intervals than during all three SF intervals, indicating lower HR during the PVT, which was the last task of the experiment. There was also a significant difference between all SF and PVT intervals in the pairwise comparison. These results suggest that it may be feasible to use both HR and IBI to detect increased fatigue across different task types, for a shorter period of fatigue (25 minutes), while the same measures might not detect fatigue within the same task. Furthermore, both duration and task type are critical factors when monitoring and detecting changes in mental states.

### SBP and DBP

Although the use of BP in detecting mental state changes is not as common as the use of HR or HRV, prior work has linked BP to changes in workload, stress, and fatigue [18]. In line with prior work, we expected both SBP and DBP to be sensitive to differentiating between tasks within the workload, stress, and fatigue conditions. Both SBP and DBP could significantly differentiate between math and preparation for TSST (*t >* 2), before but not after correcting for multiple comparisons (*p* > .05). Other models yielded nonsignificant results for all tasks.

Both SBP and DBP have been found to be sensitive measures of increased cognitive workload [9]. In a study by Hjortskov et al. [9] continuous BP measures revealed that both SBP and DBP could significantly differentiate all the conditions from the three resting periods. However, when it came to differentiating between the different stress sessions (and control) SBP did not distinguish between them, and DBP actually increased during the control condition compared to the stress conditions. Sequeira et al. [15] used BP to indicate changes in cardiovascular reactivity during three different stress conditions (TSST, singing task, and unsolvable anagrams) and found that only the TSST elevated BP. Both studies set their significance level at *p* = .05, but exact significance values were not reported. Although not a direct measure of sensitivity, *p-*values can provide some indication of how well a cardiovascular measure differentiates between tasks. This underscores the need to report exact test statistics (e.g., t, p) that indicate how well cardiovascular measures differentiate between tasks.

### HRV-RMSSD

HRV-RMSSD was another cardiovascular variable we expected to differentiate among tasks across the three conditions: cognitive workload, stress, and fatigue. The cognitive model yielded significant results between tasks, whereas no statistical differences were found between the TSST conditions. The fatigue model showed significant differences between tasks, but none were significant after correcting for multiple comparisons. The pairwise comparison for our cognitive model showed that RMSSD could differentiate between BART and Stroop. However, and more interestingly, RMSSD was able to differentiate between CORSI and Stroop, and JLAP and Stroop. The results were highly significant, indicating that RMSSD was sensitive to changes between the Stroop and the other three cognitive tasks. It is important to note that, of the four cognitive tasks, the Stroop task may be the one that elicits not only workload but also stress. It is therefore possible that HR and IBI, which distinguished BART from the other tests, are better suited to distinguishing faster-paced reaction tests that rely on working memory and attention from tasks that rely more on strategy and long-term knowledge. RMSSD may, however, be more sensitive to detect differences in stress levels.

Prior work has shown that RMSSD is not always sensitive to detecting different levels of workload [3,18]. For example, in their study on emergency physicians in Germany, previously discussed, five-minute segments of RMSSD were only sensitive to indicate increased workload during primary care time compared with baseline, but not during the drive to the scene [3]. Similarly, in a driving simulation study using three versions of the n-back task (0-back, 1-back, and 2-back) to simulate multitasking while driving, RMSSD measured over two-minute segments differentiated between low and higher workload levels, but not between medium and high workload [18]. In contrast, other studies have found RMSSD to be sensitive to differentiate between all conditions of the studies [14,28]. However, the temporal factor alone does not seem to fully explain whether RMSSD can differentiate between less robust changes in workload.

Another study, also using a driving task and two-minute segments of RMSSD while participants performed cognitive tasks with low and high n-back demands compared with baseline, found RMSSD to be quite sensitive to even small changes in increased cognitive workload [28]. As previously stated, we also expected RMSSD to differentiate among the TSST conditions. The results showed that RMSSD could not differentiate between any of the conditions (preparation, speech, and math). These results were partly surprising, given RMSSD’s ability to detect autonomic nervous system activity, particularly the parasympathetic branch, thereby indirectly providing information about sympathetic nervous system activity [59,60]. We speculate that one possible explanation for these results might be floor effect; since participants were already in the TSST, widely acknowledged to decrease RMSSD [106–109], perhaps further decrease was generally more difficult since the PNS may already have been near a physiological floor in terms of the stressor. RMSSD has been shown to distinguish between restful periods and public speaking [56,57]. But here, the focus was on difference scores and to what extent the metric could distinguish between smaller changes in stress (between the different TSST conditions). Some of the studies that have shown a decrease in RMSSD have compared the TSST conditions to baseline, which would, in general (and also in the current study), show a greater difference between the means rather than between the different already stressful conditions, making it, in general, more likely to get a statistical difference. Still, partly to our surprise, the current study indicates that other cardiovascular variables, such as HR and IBI, might be better at capturing small changes in cardiovascular reactivity during TSST than RMSSD.

Finally, we expected RMSSD to be sensitive to detect small changes in fatigue over time. Although the model showed significant differences between the third and last interval of the PVT, as well as across all three intervals of SF, compared to the reference task PVT interval one, these results did not remain significant after correction for multiple comparisons. After correcting for multiple comparisons, there was only a so-called marginal significance (*p* = .060) between PVT part one and PVT part three, as well as SF part two (*p* = .0501) and SF part three (*p =* .054), indicating a trend toward lower RMSSD during those intervals compared to PVT part one. These results are partly surprising, since increased RMSSD could be interpreted as increased fatigue, given the association between RMSSD and the parasympathetic nervous system [59,60]. At least one study has previously associated increased RMSSD with fatigue [17]. However, results regarding HRV in general and fatigue have been somewhat mixed [13].

### High-frequency power

Finally, we expected HF power to be sensitive enough to differentiate between tasks in the cognitive workload, stress, and fatigue conditions. No significant results were found for the fatigue or stress model. These results do not align with some of the studies reviewed for the current paper. For example, one study found HF to be able to differentiate between the control condition and stress condition, as well as three resting periods [9]. Still, the same study was unable to differentiate between baseline and stress. Another study also concluded that mental stress was associated with lower HF [110]. However, the designs of those two studies were very different from those of the current study. In addition, the current study found no differences between the fatigue conditions. In contrast, the cognitive-workload model showed significant pairwise differences between Corsi and Stroop and between JLAP and Stroop. These results align with the literature on HF and cognitive workload. For instance, a study by Mehler et al. [28] found significant differences in HF power between the difficult version of the n-back and baseline, but not between the easier version of the n-back and baseline. Those results indicate that the type of cognitive workload matters, which is consistent with our findings.

Another factor that might be of importance is the temporal factor; studies that have found differences between HF power and increased workload have varied in the duration of the ECG segments used to calculate HRV. For instance, in their study on emergency physicians, Schneider et al. [3] used five-minute segments and were able to differentiate between baseline and driving to the scene, as well as baseline and attending to the injured or sick at the scene. Suggesting that HF was able to detect more subtle changes in workload than RMSSD. Mehler et al. [28] reported similar results in their study, using two-minute segments of HF power while participants drove and performed low-and high-workload versions of the n-back task. HF power could differentiate between high workload and baseline, but not between low workload and baseline.

In contrast, two-minute segments of HF power were unable to differentiate between low, medium, or high n-back tasks meant to resemble multitasking while driving in a study by Tjolleng et al. [18]. Three-minute segments of HF power have also been reported to differentiate between both workload and stress periods [9] while segments as long as 15 minutes have been reported to identify fatigue over time [17]. These results are relevant to the current study, which used segments of varying lengths to measure HRV, mostly due to differences in task durations. In line with the recommendations for HRV measures [48,49] we sought to use five minutes where possible, but we used the whole task when the task did not reach five minutes. The length of the HRV recordings, therefore, ranged from a minimum of one minute to a maximum of 5 minutes. Although not always directly assessing fatigue and stress, given the findings above regarding different durations of HRV inspection, we do not conclude that the temporal factor alone explains why HF power was not sensitive to detecting stress and fatigue.

### Summary and recommendation

The current study found HR and IBI to be generally more sensitive measures for detecting changes in the following mental states: cognitive workload, acute stress and time-on-task fatigue compared with IBI derived HRV measures (RMSSD and HF) and BP. Moreover, the results confirmed previous findings [21,22] suggesting that the sensitivity of cardiovascular measures to mental state changes is highly dependent on task type and the relevant cognitive processes required for the task. Both HR and IBI are sensitive to changes in acute high stress (public speaking etc.) but their sensitivity varies with duration and type of task for fatigue detection. For cognitive workload, these measures are highly sensitive to tasks relying more on attentional and working memory processes rather than those linked with long-term memory and strategy where changes in workload are slow and perhaps cumulative over time. BP did not appear to be a sensitive measure in the current study for mental state changes. Both RMSSD and HF should be used along with HR and possibly IBI. In the current study RMSSD was sensitive to the Stroop test compared with the other cognitive tasks possibly indicating that the measure is sensitive to detect cognitive workload linked with stressful, fast-paced tasks.

Prior work has demonstrated that both the type and length of a task can have an impact on whether, or to what extent, cardiovascular reactivity changes [21,22]. This partly reflects the temporal characteristics of the metrics themselves: HR can detect rapid, moment-to-moment changes in physiological activation, whereas HRV is typically calculated over longer time windows and may therefore capture regulatory dynamics over slightly extended periods [49]. For example, reaction-time tasks may induce different HR and HRV responses compared to those requiring sustained working memory or problem-solving skills, possibly due to distinct underlying cognitive (brain) processes [21,22]. Different cognitive processes, likely activating distinct brain regions, yield varied physiological outputs, underscoring the importance of considering task type in cardiovascular response studies. Future studies on mental state monitoring should be mindful of using a varied task design that includes multiple cognitive challenges and intensities to better understand how these cardiovascular measures relate to dynamic changes in mental states.

### Limitations

For the current study, several limitations must be considered. Firstly, the current study did not control for mental disorders such as depression, anxiety, PTSD, and OCD, all known to affect cardiovascular activity [113–120]. Secondly, the current study did not control for medications and caffeine consumption, which are also believed to affect cardiovascular activity [120,121]. Thirdly, baseline measurements were taken in the same room as the experiment just before participants underwent the experiment. Therefore, it cannot be ruled out that anticipatory stress affected the baseline measurements, thereby influencing the comparison between baseline and cognitive tests and the TSST. Another limitation worth considering is that missingness in the data due to noise or other technical issues resulted in uneven sample sizes. Uneven sample sizes across models might have influenced our results; however, the mean age, its standard deviation, and gender ratio were usually remarkably similar. Still, it cannot be overlooked that other confounding factors associated with the varying sample size described above might have influenced the results. The final limitation is that, due to noise and technical issues in the Stroop data, only the first 12 minutes of the 15-minute task were used for analysis; therefore, the segment analyzed might not have fully captured changes in cardiovascular reactivity during SF.

## Conclusion

Studies on cardiovascular reactivity are a promising approach for indicating changes in cognitive workload, stress, and fatigue [9,13,15,20,28]. The current study demonstrated that cardiovascular variables, especially those that rely on HR, were sensitive in differentiating between several cognitive tasks and the TSST. However, we also demonstrated challenges with cardiovascular measures, such as differentiating between JLAP and Corsi, and detecting changes in cardiovascular reactivity over time within the same task, especially SF and PVT. In addition, SBP and DBP were not sensitive to any of the conditions after correcting for multiple comparisons. Future studies should aim to include more cognitive tests and various stressors to clarify the relationship between cardiovascular reactivity and mental states, as a limitation of the current literature is that many studies use only one cognitive test across different levels. The high correlation between shorter and longer HRV durations may limit its ability to detect changes in fatigue over time in such prolonged tasks. Therefore, we recommend further investigation into whether cardiovascular changes detect increased fatigue during prolonged tasks, e.g., even longer Stroop tasks.

## Supporting information

S1 Code. R code used to reproduce findings of the current study

S2 Dataset. The dataset underlying the findings of the current study is provided in RSD format

S3 Tables (S1-S18). Complete results from all regression models and pairwise comparisons

S4 The dataset underlying the findings of the current study is provided in CSV format

## Notes

### Competing Interest Statement

The authors have declared no competing interest.

